# Mutation in the matricellular gene fibulin-4 leads to endothelial dysfunction in resistance arteries

**DOI:** 10.1101/2022.05.20.492867

**Authors:** Michelle Lin, Kara Jones, Bridget M. Brengle, Robert P. Mecham, Carmen M. Halabi

**Author notes:** Corresponding author: Carmen M. Halabi, 660 South Euclid Ave, Campus Box 8116, Saint Louis, MO 63110.

## Abstract

Mutations in fibulin-4 (*FBLN4*), a matricellular gene required for extracellular matrix (ECM) assembly, result in autosomal recessive cutis laxa type 1B (ARCL1B), a syndrome characterized by loose skin, aortic aneurysms, pulmonary emphysema and skeletal abnormalities.

*Fbln4*^*E57K/E57K*^ mice recapitulated the phenotypes observed in ARCL1B. In particular, they exhibited ascending aortic aneurysms, elastic fiber fragmentation and increased stiffness in large arteries, and systolic hypertension. Surprisingly however, internal elastic laminae of small resistance and muscular arteries were intact. Here, we show that the increased pulsatile flow resulting from the structural abnormalities and increased stiffness of conduit arteries in *Fbln4*^*E57K/E57K*^ mice leads to increased shear stress, a highly oxidative environment, and endothelial dysfunction related to reduced nitric oxide bioavailability in resistance mesenteric arteries. These data have significant implications, not only for the basic biology of ECM assembly along the arterial tree, but also for the clinical consequences of large artery stiffness on the microcirculation.

## Introduction

Arteries fulfill differing roles along the arterial tree. Large proximal arteries serve as conduits that propel blood forward and dampen the pulse pressure generated by each heartbeat, allowing for laminar blood flow distally^1^. This pulse dampening property of large arteries, termed the Windkessel effect, is afforded by the protein elastin, which accounts for 40-60% of the aorta’s dry weight depending on species^2^. Distal muscular and resistance arteries on the other hand provide the vascular tone necessary for tissue perfusion and have little elastin. Abnormal arterial elastin assembly or its fragmentation, as occurs with aging and when accelerated in certain disease states such as aneurysms, diabetes and chronic kidney disease^3-9^, results in large artery stiffness, a now well-recognized independent risk factor for cardiovascular mortality^10-15^.

Elastic fiber assembly is a highly orchestrated process that involves interactions between numerous proteins and extensive crosslinking of the monomer tropoelastin by the enzyme lysyl oxidase (LOX) to generate the highly stable polymer, elastin with a half-life of over 70 years^16,17^. One molecule that plays a critical role in elastin assembly is fibulin-4 (FBLN4). FBLN4, also known as epidermal growth factor-containing fibulin-like extracellular matrix protein 2 (EFEMP2), belongs to an eight-member family of extracellular matrix proteins that share significant structural homology characterized by repeats of calcium-binding epidermal growth factor-like (cbEGF) domains and a carboxy-terminus fibulin domain^18,19^. While their modular structure is similar, fibulins are involved in varied cellular functions including cell growth, adhesion, and motility as well as the formation and stability of basement membranes, elastic fibers and connective tissues^20-22^. FBLN4, in particular, plays a critical role in elastic fiber assembly as mice lacking the gene die perinatally secondary to aortic aneurysm rupture, pulmonary emphysema and diaphragmatic hernias due to the absence of intact elastic fibers^23^. Furthermore, humans with homozygous or compound heterozygous mutations in *FBLN4* develop autosomal recessive cutis laxa type 1B (ARCL1B), a syndrome characterized by loose inelastic skin, ascending aortic aneurysms, pulmonary emphysema, joint laxity and skeletal deformities, with underdeveloped elastic fibers on skin biopsy^24-26^.

A mouse model carrying a disease-causing mutation in *Fbln4* (E57K) has been generated^27^. This mutation results in a glutamate to lysine change at position 57 (E57K), a highly conserved amino acid residue in the first modified cbEGF domain of FBLN4. Unlike global *Fbln4* deletion, which led to perinatal lethality^23^, mice with mutant FBLN4 survived up to two years of age^27,28^, suggesting that the mutation is not a complete loss-of-function mutation. *Fbln4*^*E57K/E57K*^ mice were shown to recapitulate the phenotypes seen in ARCL1B^27^. Specifically, mutant mice developed the characteristic ascending aortic aneurysms, emphysema and skeletal abnormalities related to abnormalities in elastin and collagen fibers. To further delineate the cardiovascular consequences of mutant FBLN4, we performed structural and functional characterization of *Fbln4*^*E57K/E57K*^ mice and found that they developed significant large artery stiffness and systolic hypertension^28^. Surprisingly however, while large arteries exhibited elastic fiber fragmentation and vessel wall disarray including changes in smooth muscle cell morphology and an increase in medial thickness, resistance arteries including renal, saphenous and mesenteric arteries were structurally intact when examined by transmission electron microscopy^28^. These data suggested that elastin assembly may have different molecular requirements based on vessel type.

Given the crosstalk between large and resistance arteries, the significant contribution of resistance arteries to blood pressure regulation and the potential change in smooth muscle cell phenotype due to mutant FBLN4^29,30^, we sought to determine the functional consequences of mutant FBLN4 on resistance arteries. Here, we show that *Fbln4*^*E57K/E57K*^ mesenteric arteries retained the ability to constrict to pressure and vasoactive substances, but exhibited impaired endothelial-dependent vasodilation. Furthermore, we show that the endothelial dysfunction in *Fbln4*^*E57K/E57K*^ mesenteric arteries is secondary to impaired nitric oxide (NO)-mediated vasodilation caused by increased oxidative stress and decreased endothelial nitric oxide synthase (eNOS) activation. Lastly, we identified changes in endothelial cell polarity in *Fbln4*^*E57K/E57K*^ mesenteric arteries, indicating increased shear stress. These data emphasize the importance of targeting elastin fragmentation and large artery stiffness to avoid its negative consequences on the microcirculation, which has broad implications not only for patients with cutis laxa, but potentially for all conditions associated with elastic fiber fragmentation and large artery stiffness.

## Results

### *Fbln4*^*E57K/E57K*^ mesenteric arteries have preserved myogenic tone and contractile ability

Unlike conduit arteries which function as elastic reservoirs to reduce pulsatile pressure allowing laminar blood flow distally, resistance or muscular arteries function to regulate peripheral vascular tone and maintain constant blood flow to support end organ perfusion. This ability of resistance arteries to maintain constant blood flow is conferred, at least in part, by the myogenic tone^31^. Although the *Fbln4*^*E57K/E57K*^ resistance arteries were unaffected ultrastructurally^28^, given the systolic hypertension and elevated pulse pressure noted in *Fbln4*^*E57K/E57K*^ mice and FBLN4’s role in mediating ECM assembly, we sought to determine whether *Fbln4*^*E57K/E57K*^ resistance arteries exhibited any functional impairment. We did so by assessing the passive (in the absence of calcium) and active (in the presence of calcium) diameters of *Fbln4*^*E57K/E57K*^ and littermate control mesenteric arteries in response to increasing pressure. As shown in figure 1A, both passive and active diameters of *Fbln4*^*E57K/E57K*^ mesenteric arteries were similar to those of littermate control vessels at all pressures tested. Myogenic tone, or the ability of smooth muscle cells (SMCs) to constrict in response to increasing pressure, was unaffected by the mutation (Fig 1B). Similar to the vessels’ responsiveness to pressure, the ability of *Fbln4*^*E57K/E57K*^ resistance arteries to constrict to the vasoactive substances angiotensin II and phenylephrine was similar to that of littermate controls (Fig 1C&D). These data suggest that SMCs’ ability to constrict to mechanical and chemical cues is preserved in *Fbln4*^*E57K/E57K*^ mice.

**Figure 1.**
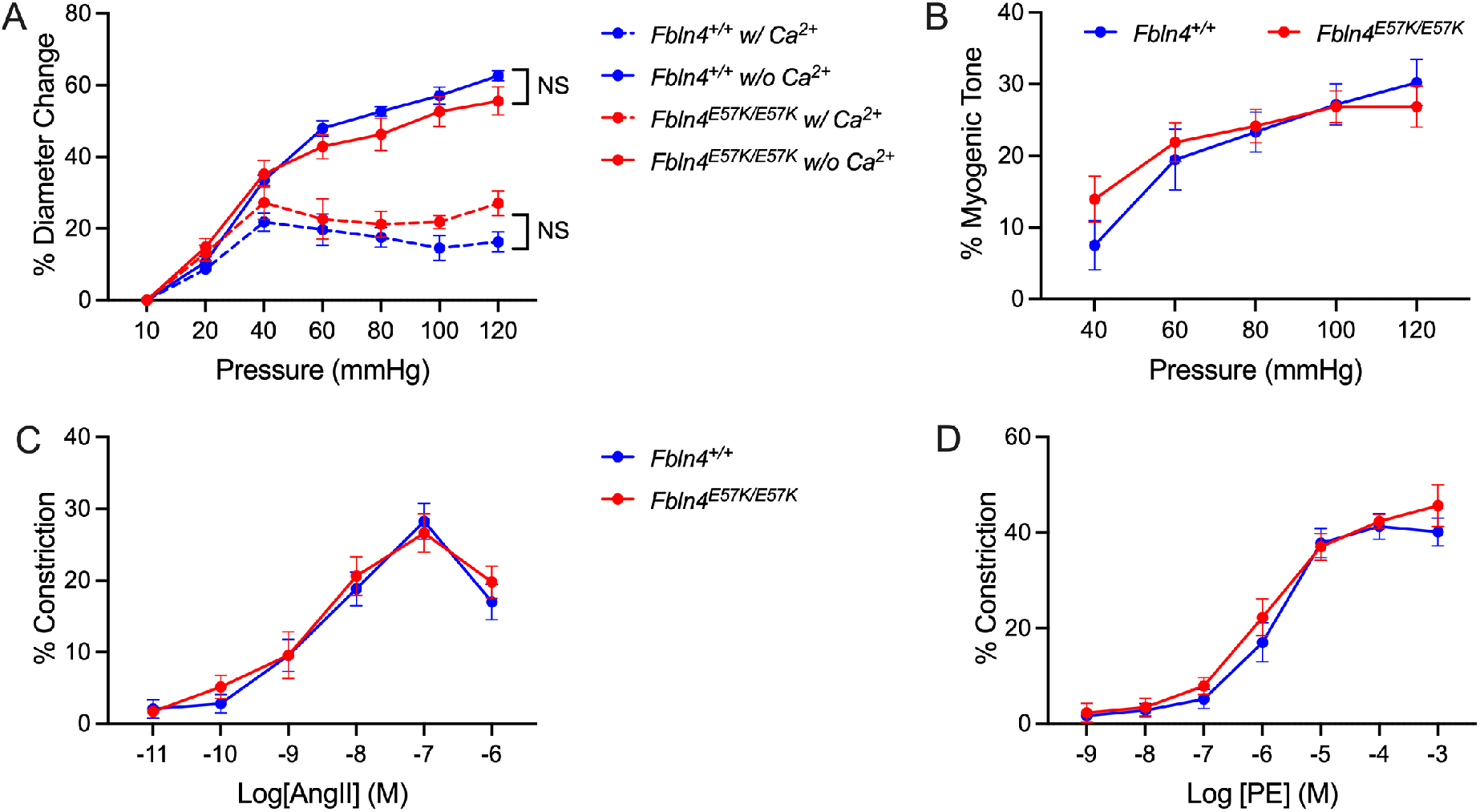
Mutation in Fibulin-4 does not affect mesenteric artery myogenic or contractile responses. (A) Pressure-diameter relationships of third order mesenteric arteries isolated form *Fbln4*^*E57K/E57K*^ (n=10) and littermate control (n=9) mice reflecting active (dashed lines) and passive (solid lines) diameter changes in the presence and absence of calcium, respectively. (B) % myogenic tone of third order *Fbln4*^*E57K/E57K*^ and littermate control mesenteric arteries calculated as (passive diameter – active diameter)/passive diameter * 100. (C&D) Dose-response curves showing % constriction of third order *Fbln4*^*E57K/E57K*^ (n=7) and littermate control (n=6) mesenteric arteries to increasing doses of angiotensin II (angII, C) and phenylephrine (PE, D). Data are presented as mean ± standard error of the mean. Two-way analysis of variance with Tukey’s multiple comparison test was performed to compare all groups. NS = no significant difference.

### *Fbln4*^*E57K/E57K*^ mesenteric arteries display endothelial dysfunction that is nitric oxide-dependent

The presence of systolic hypertension and increased pulsatile pressure often leads to impaired vasodilation of resistance arteries, further exacerbating the hypertension^32^. We examined the ability of *Fbln4*^*E57K/E57K*^ mesenteric arteries to respond to the endothelium-dependent and endothelium-independent vasodilators acetylcholine and papaverine, respectively. As shown in figure 2A&B, while *Fbln4*^*E57K/E57K*^ mesenteric arteries dilated maximally to papaverine, they had an impaired response to acetylcholine compared to littermate control vessels, suggesting endothelial dysfunction.

**Figure 2.**
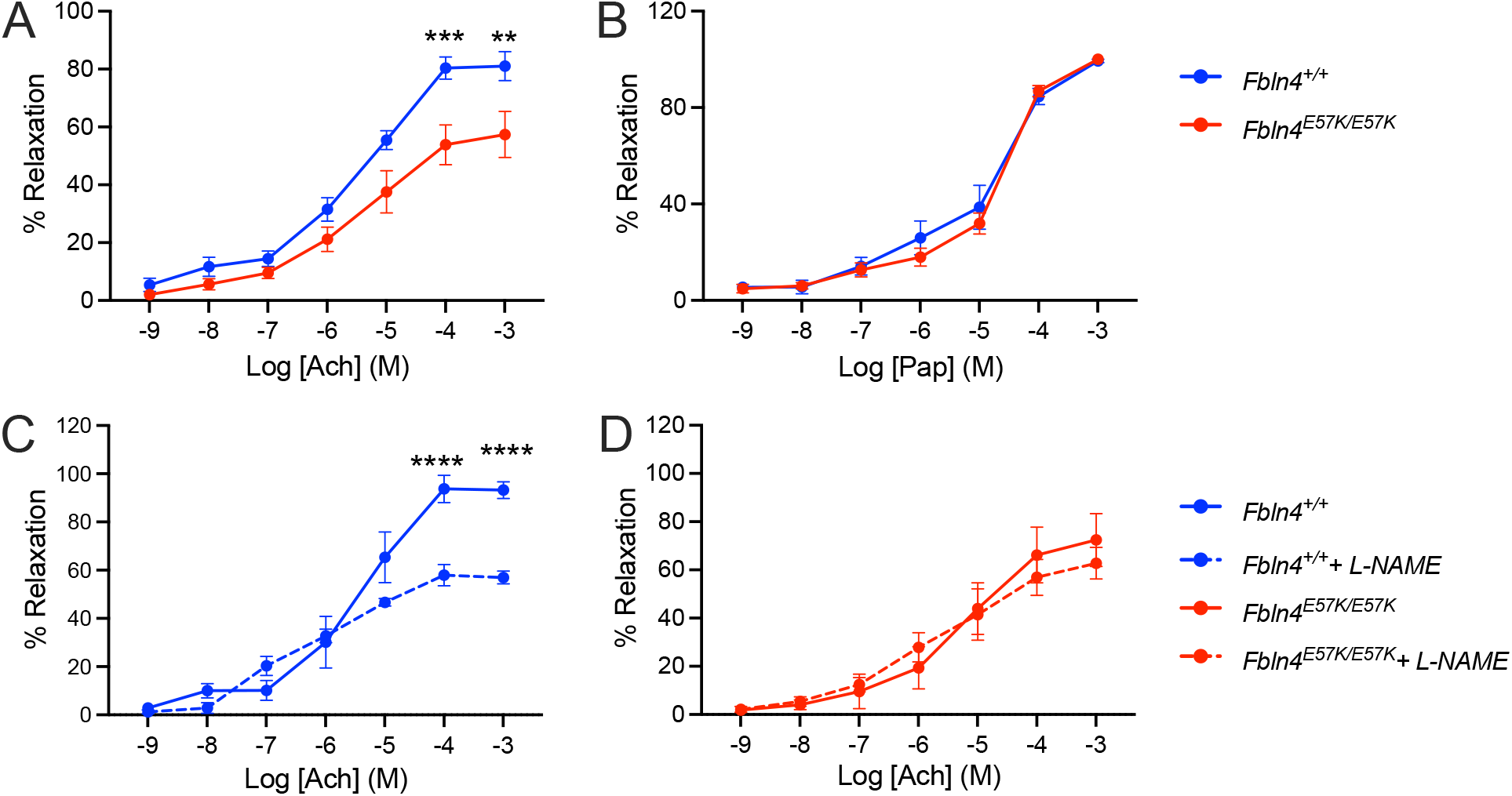
*Fbln4*^*E57K/E57K*^ mesenteric arteries have impaired endothelial-dependent vasodilation that is nitric oxide-mediated. (A&B) Dose-response curves showing % relaxation of third order *Fbln4*^*E57K/E57K*^ (n=7) and littermate control (n=6) mesenteric arteries to increasing doses of acetylcholine (Ach, A) and papaverine (Pap, B). (C&D) Dose-response curves showing % relaxation of wild-type (n=7, C) and *Fbln4*^*E57K/E57K*^ (n=8, D) third-order mesenteric arteries to increasing doses of acetylcholine (Ach, A) in the presence (dashed lines) or absence (solid lines) of the eNOS inhibitor, L-NAME. Data are presented as mean ± standard error of the mean and compared using two-way analysis of variance with Sidak’s multiple comparisons test. ** P<0.005, *** P<0.001, **** P<0.0001

Endothelial-dependent vasodilation is mediated via nitric oxide, endothelium-derived hyperpolarizing factor and prostacyclins^33,34^. To determine whether, nitric oxide-mediated dilation is affected in *Fbln4*^*E57K/E57K*^ mesenteric arteries, we examined the vessels’ responsiveness to acetylcholine in the presence of the endothelial nitric oxide synthase (eNOS) inhibitor, L-N^G^-Nitro arginine methyl ester (L-NAME). As shown in figure 2C&D, eNOS inhibition led to a significant reduction in vasodilation in response to acetylcholine in wild-type (WT, Fig2C), but not *Fbln4*^*E57K/E57K*^ mesenteric arteries (Fig 2D), indicating decreased NO bioavailability in mutant vessels.

### eNOS expression is unaffected, but its activation is abrogated in *Fbln4*^*E57K/E57K*^ mesenteric arteries

To determine whether the decreased NO bioavailability in *Fbln4*^*E57K/E57K*^ mesenteric arteries is due to decreased NO production, we first assessed expression of the three NOS isoforms (nNOS, iNOS and eNOS) in *Fbln4*^*E57K/E57K*^ and littermate control mesenteric arteries. As shown in figure 3A, expression of all NOS isoforms in *Fbln4*^*E57K/E57K*^ mesenteric arteries was similar to that of littermate control vessels. eNOS is the major NOS isoform in the vasculature and its posttranslational modification by phosphorylation or dephosphorylation at different sites alters its enzymatic activity^35,36^. For instance, phosphorylation of serine 1177 leads to increased eNOS activity while phosphorylation of threonine 495 inhibits its activity^37^. We examined eNOS phosphorylation at Serine 1177 in *Fbln4*^*E57K/E57K*^ and control mesenteric arteries by western blotting and found reduced eNOS phosphorylation (Fig 3B), suggesting reduced activity in mutant vessels.

**Figure 3.**
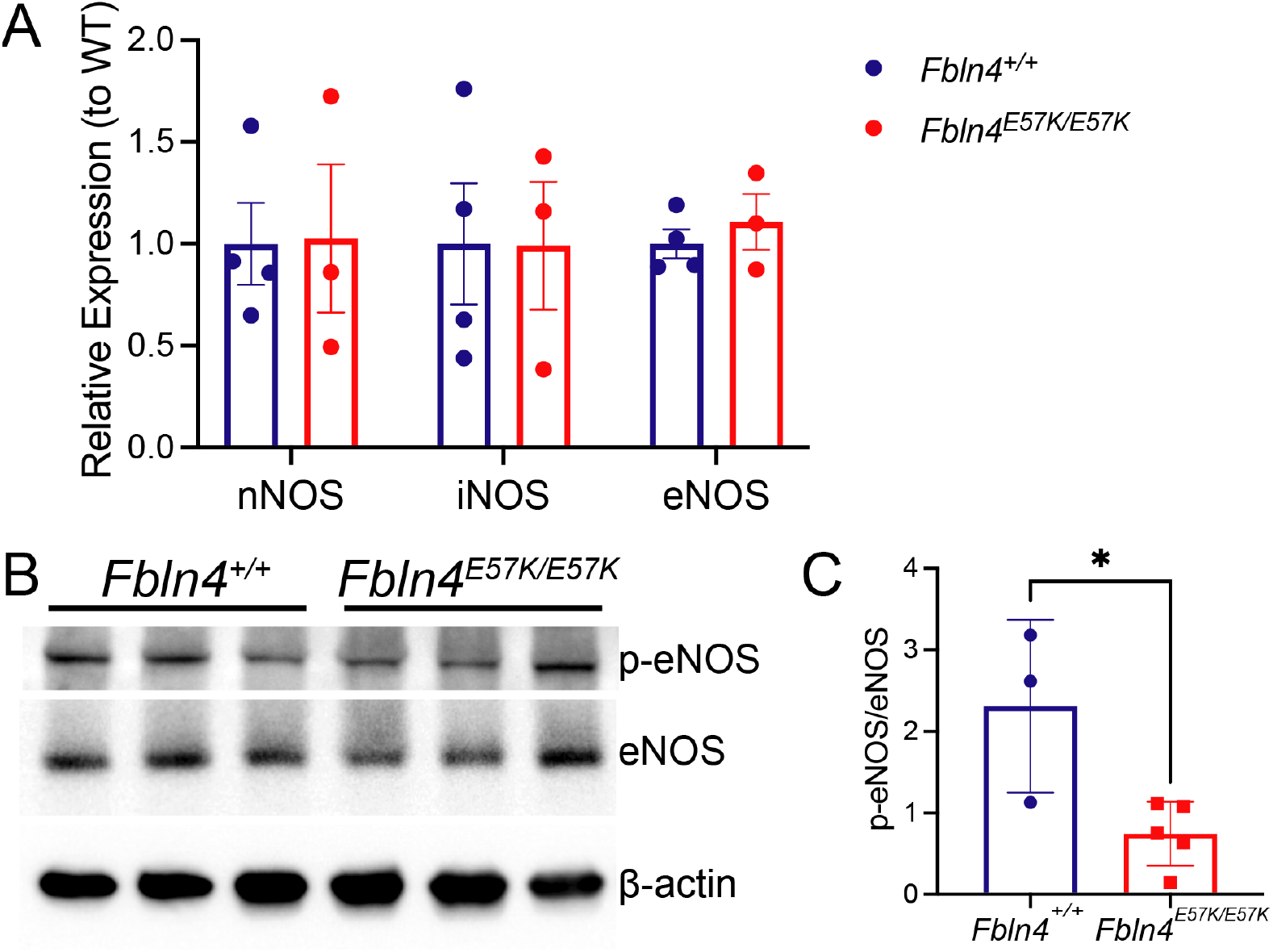
While eNOS expression is unaffected, its phosphorylation is decreased in *Fbln4*^*E57K/E57K*^ mesenteric arteries. (A) Gene expression of the three NOS isoforms in *Fbln4*^*E57K/E57K*^ and littermate control mesenteric arteries by qRT-PCR. Gene expression was normalized to that of β2-microglobulin and presented relative to the average gene expression in WT vessels. Data are presented as mean ± SEM and compared using two-way analysis of variance with Sidak’s multiple comparisons test (B) Western blots of mesenteric artery lysates from *Fbln4*^*E57K/E57K*^ and littermate control mice using S1177-p-eNOS, eNOS and β-actin antibodies. (C) Quantification of the Western blot data comparing p-eNOS to total eNOS levels in *Fbln4*^*E57K/E57K*^ and littermate control mesenteric arteries. Data are presented as mean ± SD and compared using student’s t-test. * P<0.05.

### *Fbln4*^*E57K*^ mesenteric arteries display increased oxidative stress

To catalyze the production of NO, eNOS needs to form a homodimer in order to bind the necessary co-factor tetrahydrobiopterin (BH_4_) and the substrate L-arginine^38^. As a monomer, eNOS generates superoxide (O_2_^-^) rather than NO from its oxygenase domain, a process termed eNOS uncoupling^35^. Additionally, in the presence of oxidative stress, BH4 is oxidized to the BH_2_ leading to eNOS uncoupling and further production of O_2_^-39^. Using the superoxide indicator, dihydroethidium (DHE), we assessed superoxide levels in mesenteric arteries of *Fbln4*^*E57K/E57K*^ and littermate control mesenteric arteries. As shown in figure 4, *Fbln4*^*E57K/E57K*^ mesenteric arteries exhibit a significant increase in DHE staining compared to littermate control vessels, suggesting increased oxidative stress in mutant vessels.

**Figure 4.**
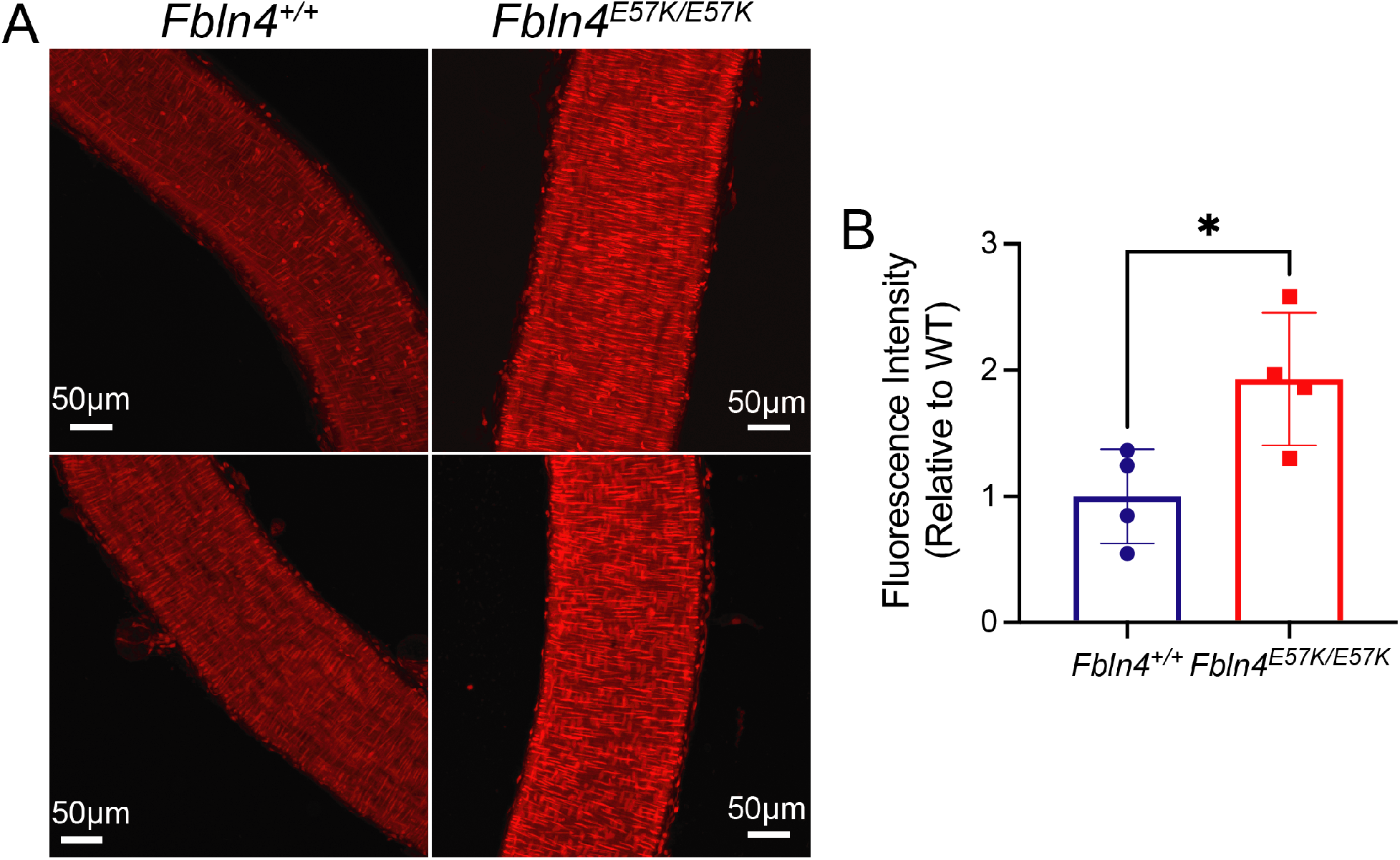
*Fbln4*^*E57K/E57K*^ mesenteric arteries exhibit increased superoxide production. (A) Maximum intensity images of z-stacks of *Fbln4*^*E57K/E57K*^ and littermate control third order mesenteric arteries stained with dihydroethidium (DHE) and obtained using confocal microscopy. (B) Quantification of the fluorescence intensity of DHE-stained vessels shown in A, using the average of three regions of interest per mouse and shown relative to the average fluorescence intensity of WT vessels. Data are presented as mean ± SD and compared using student’s t-test. * P<0.05.

### *Fbln4*^*E57K/E57K*^ mesenteric arteries exhibit alterations in endothelial cell polarity and internal elastic lamina fenestration number

Given the increased large artery stiffness and pulse pressure in *Fbln4*^*E57K/E57K*^ mice^28^, we posited that the increased pulsatile flow would result in increased hemodynamic shear stress in the microcirculation. To assess changes in shear stress, we determined the proportion of cells with Golgi orientation with, against or at an angle to blood flow. At physiologic shear stress level, the endothelial cell orientation is parallel to blood flow and the Golgi orientation is generally upstream of the nucleus (i.e. against the flow direction)^40,41^. As shown in figure 5A&B, the proportion of cells with the Golgi pointing against flow was increased in endothelial cells of *Fbln4*^*E57K/E57K*^ mesenteric arteries compared to WT littermate vessels, indicating increased shear stress. In addition to affecting cell polarity, hemodynamic forces have been shown to affect internal elastic fenestrae density^42,43^. We examined fenestrae number in the internal elastic laminae (IEL) of *Fbln4*^*E57K/E57K*^ and littermate control mesenteric arteries and found a significantly greater number of fenestrae in IEL of mutant vessels (Fig 5C&D).

**Figure 5.**
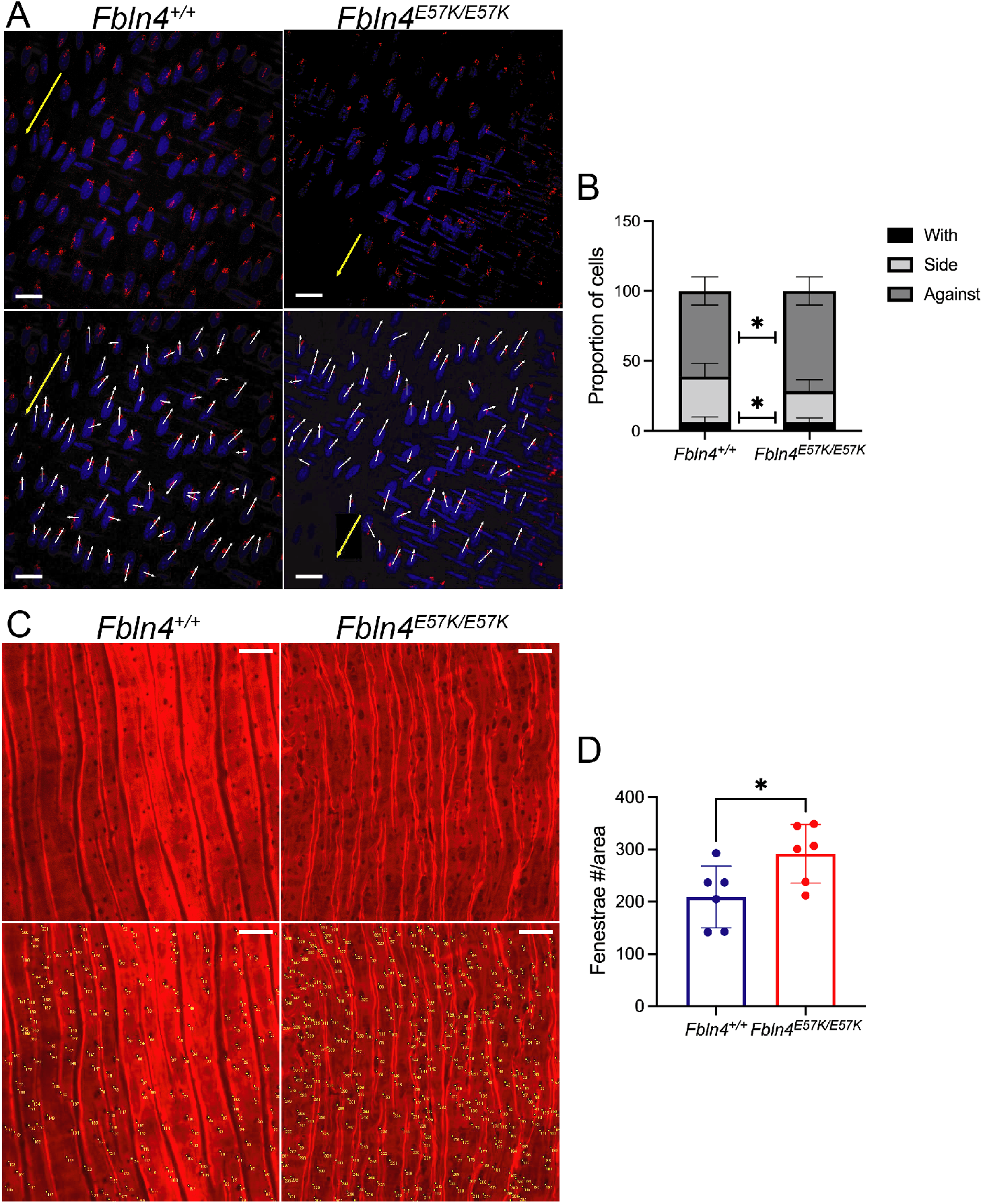
*Fbln4*^*E57K/E57K*^ mesenteric arteries exhibit alterations in endothelial cell polarity and IEL fenestration number. (A) Representative maximum intensity images of *en face* WT and *Fbln4*^*E57K/E57K*^ third order mesenteric arteries immunostained with Golgi antibody (GM130, red) and DAPI (blue) to determine Golgi orientation (white arrows) relative to the nucleus and blood flow (yellow arrows). (B) Quantification of Golgi position around the nucleus compared to blood flow. N = 3 mice (349 cells from 8 vessel segments) for WT and 4 mice for *Fbln4*^*E57K/E57K*^ (431 cells from 10 vessel segments). Data are presented as mean ± SD and compared using two-way analysis of variance with Sidak’s multiple comparisons test. * P<0.05. (C) Representative maximum intensity images of *en face* WT and *Fbln4*^*E57K/E57K*^ third order mesenteric arteries stained with Alexa Fluor-633 hydrazide. Yellow numbers indicate fenestrae counted. (D) Quantification of fenestrae numbers counted in C. Data are presented as mean ± SD and compared using student’s t-test. * P<0.05. Scale bars in A&C = 20 μm.

## Discussion

Using a mouse model carrying a disease-causing mutation in the matricellular gene *Fbln4*, we show that mutant mesenteric resistance arteries, while seemingly structurally intact by transmission electron microscopy, exhibit significant endothelial dysfunction, but preserved smooth muscle cell function. This endothelial dysfunction is mediated by reduced NO bioavailability, as inhibition of eNOS with L-NAME did not alter mutant mesenteric arteries’ response to acetylcholine. The data presented herein support our working model (figure 6) where increased pulsatile flow due to elastic fiber fragmentation and increased stiffness of large arteries leads to increased pulsatile shear stress in the microcirculation, and resultant increase in oxidative stress in resistance arteries. A highly oxidative environment reduces NO bioavailability via both increased consumption (NO + O_2_^-^ generates peroxynitrite, which acts a vasoconstrictor) as well as reduced production of NO (uncoupling and decreased phosphorylation of Ser1177 of eNOS) resulting in endothelial dysfunction and worsening of the hypertensive phenotype.

**Figure 6.**
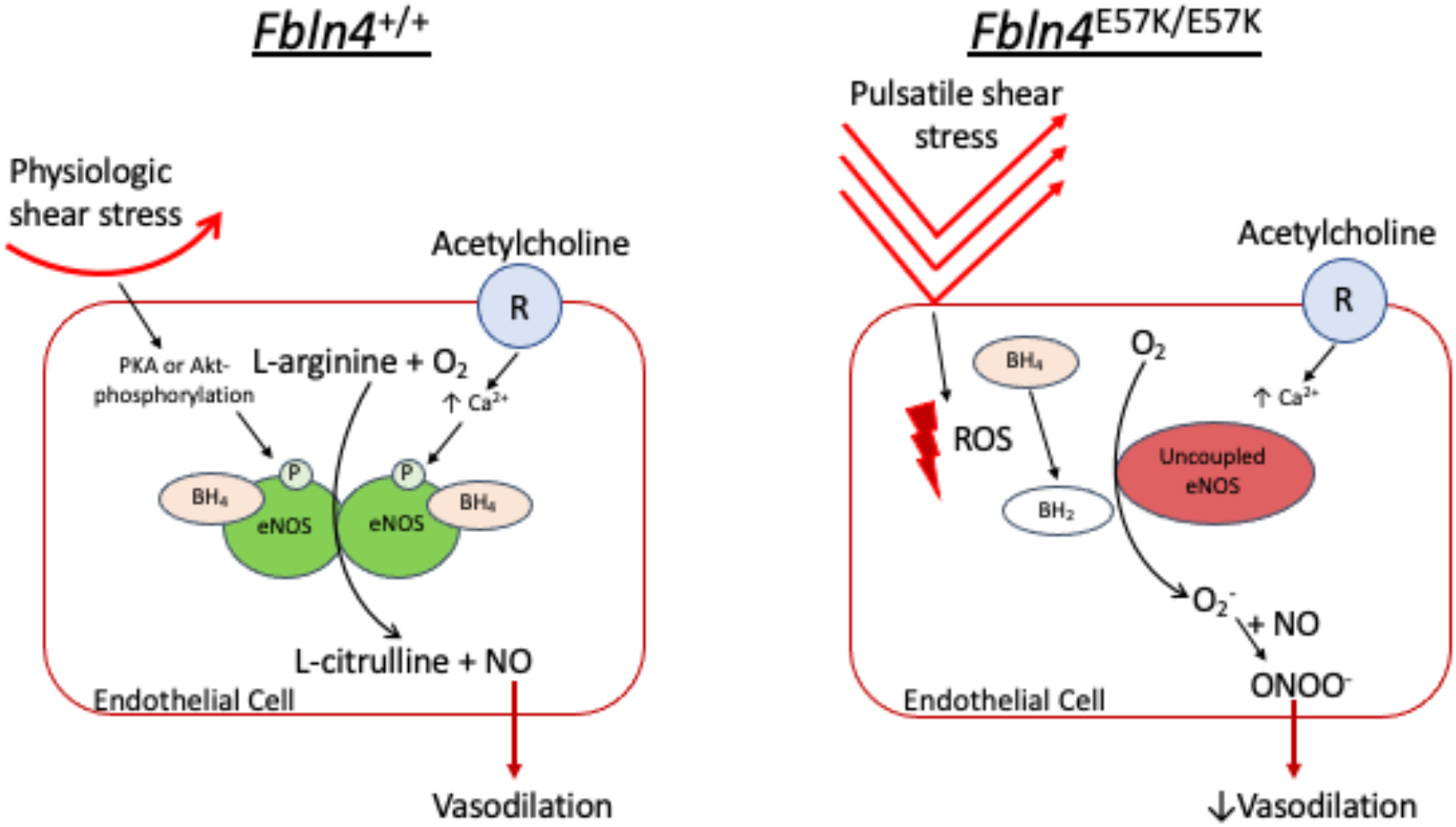
Working Model. Under physiologic conditions, either via a calcium-dependent mechanism such as acetylcholine binding to its receptor or a calcium-independent mechanism as with physiologic shear stress, eNOS dimerizes and gets activated resulting in the generation of NO, which leads to vasodilation. In *Fbln4*^*E57K/E57K*^ mice however, the increased pulsatile flow to the microcirculation leads to increased ROS production, which oxidizes tetrahydrobiopterin (BH_4_) to dihydrobiopterin (BH_2_) resulting in uncoupling of eNOS. As a monomer, eNOS generates superoxide (O_2_^-^) rather than NO further exacerbating oxidative stress. In addition to eNOS uncoupling, O_2_^-^ may interact with available NO to generate peroxynitrite (ONOO^-^), which acts as a vasoconstrictor, leading to further reduction in vasodilation in *Fbln4*^*E57K/E57K*^ resistance arteries.

The increase in oxidative stress in *Fbln4*^*E57K/E57K*^ compared to littermate control mesenteric arteries as seen by DHE staining was quite striking and is likely related to increased pulsatile shear stress as well as uncoupling of eNOS. In fact, pulsatile flow has been shown to cause reactive oxygen species (ROS) production^44,45^. An environment with increased ROS production leads to oxidation of tetrahydrobiopterin (BH_4_), a cofactor necessary for eNOS dimerization and generation of NO. In the absence of BH_4_, eNOS is uncoupled and, as a monomer, generates O_2_^-^ rather than NO, further exacerbating oxidative stress^35,37^. It is interesting to note that increased oxidative stress and endothelial dysfunction have been observed in mesenteric and cerebral arteries of elastin insufficient mice (*Eln*^*+/-*^)^46-49^. Similar to *Fbln4*^*E57K/E57K*^ mice, *Eln*^*+/-*^ mice exhibit large artery stiffness and systolic hypertension, however, this is due to reduced elastin amount as evidenced by thinning of the elastic lamellae throughout the arterial tree rather than its abnormal assembly or fragmentation^50,51^. Inhibition of NADPH oxidase 1, an enzyme responsible for ROS production, abrogated the hypertensive phenotype in *Eln*^*+/-*^ mice, suggesting that ROS contribute to blood pressure elevation in this model^49^. It is important to point out that the endothelial dysfunction was not uniformly due to reduced NO bioavailability as

L-NAME treatment had differential effects in mesenteric vs. cerebral arteries. Furthermore, while mesenteric and cerebral arteries exhibited endothelial dysfunction, skeletal muscle feed arteries did not, suggesting vascular bed-specific differences^47,48^. It will be interesting to determine whether such endothelial dysfunction is universal or vascular bed-specific in *Fbln4*^*E57K/E57K*^ mice.

Given the physiologic similarities with the *Eln*^*+/-*^ mouse model and the potential change in SMC phenotype due the *Fbln4* mutation^29^, we were surprised to see that the vasoconstrictor responses of *Fbln4*^*E57K/E57K*^ mesenteric arteries were unaltered. In particular, active mesenteric artery constriction to increasing pressure (myogenic response) and its constriction to the vasoactive substances angiotensin II and phenylephrine were not different from those of control littermate vessels. This is in contrast to the *Eln*^*+/-*^ mouse model that showed an increased contractile response to angiotensin II in both mesenteric^47^ and cerebral arteries^52^, that was partially mediated by the angiotensin II type 2 receptor^53^. These data point to changes in signaling pathways that result from reduced elastin production compared to its fragmentation, perhaps making the *Fbln4*^*E57K/E57K*^ mouse model a good one to study target organ dysfunction related to elastic fiber fragmentation and large artery stiffness as occurs with aging.

The preserved passive response (i.e. in the absence of calcium) of *Fbln4*^*E57K/E57K*^ mesenteric arteries to increasing pressure (figure 1A) is quite intriguing. On the one hand, this observation validates that the material properties of the vessel wall are not significantly different than those of a WT vessel despite the presence of an increased number of fenestrae, corroborating the previous ultrastructural observations as well as data showing unchanged desmosine (amino acid specific to elastin) and hydroxyproline (found in collagen) content in *Fbln4*^*E57K/E57K*^ mesenteric arteries^28^. On the other hand however, the fact that the ECM of *Fbln4*^*E57K/E57K*^ mesenteric arteries is not affected by the mutation raises a question about the role of FBLN4 in ECM assembly in muscular and resistance arteries. This is especially true in light of recent evidence showing that FBLN4, through incorporation of copper in the lysine tyrosyl quinone (LTQ) domain, is required for LOX activation^54^, hence elastin and collagen crosslinking, and that the E57K mutation in FBLN4 impairs its interaction with LOX and mitigates its activation. It is interesting to speculate that another LOX family member, potentially LOXL1, is compensating for decreased LOX activity. This would imply that a molecule other than FBLN4 is responsible for the transfer of copper to the LTQ domain of other LOX family members since they all require the addition of copper for enzymatic activity. Perhaps, FBLN5 serves that role for LOXL1 given that *Loxl1* and *Fbln5* knock-out mice phenocopy each other and that FBLN5 has been shown to bind LOXL1^19,55-57^, another speculation that requires further investigation.

In conclusion, using a mouse model carrying a disease-causing mutation in *Fbln4*, a gene required for ECM assembly, we show that the increased pulsatile blood flow caused by elastic fiber fragmentation and increased stiffness of large arteries leads to increased oxidative stress and consequent impairment of NO-mediated vasodilation in resistance arteries, perpetuating the hypertension. In addition to the utility of this mouse model to assess and perhaps ameliorate target organ effects related to elastic fiber fragmentation and increased large artery stiffness as seen with aging and a multitude of common diseases, the data presented herein raise important questions about the differences in the molecular requirements for ECM assembly, perhaps not only between different vascular beds, but also between different tissues/organs.

## Methods

### Mice

*Fbln4*^*E57K//E57K*^ mice were previously generated and described^27^. Briefly, *Fbln4*^*E57K//E57K*^ mice carry a knock-in mutation (G to A) in exon 4 that leads to a substitution of glutamate to lysine at amino acid 57. Mice heterozygous for the knock-in allele were bred together to maintain the mouse colony. Both male and female littermates were used for all experiments. Tail DNA was used to genotype the mice, as previously described^27^. The mice were housed under standard conditions with free access to food and water. All protocols were approved by the Animal Studies Committee of Washington University School of Medicine.

### Mesenteric Artery Myogenic Tone Measurement

Following euthanasia with CO_2_, the gut from the duodenum to the cecum was excised and placed into cold normal Krebs’ buffer (NB) containing: 120 mM NaCl, 25 mM NaHCO_3_, 4.8 mM KCl, 1.2 mM NaH_2_PO_4_, 1.2 mM MgSO_4_, 11 mM glucose and 1.8 mM CaCl_2_ (pH 7.4). Third-order mesenteric arteries (MA) were excised, mounted onto glass cannulas in the chamber of a pressure myography system (114P, Danish Myo Technology, Denmark) and tied in place with two pieces of nylon suture, as previously described^58^. To remove the endothelial cells, air bubbles were passed through the lumen of the vessel. The vessel was briefly pressurized to 60 mmHg (less than 1 minute) to ensure there were no leaks. Pressure was then returned to 10 mmHg and vessels were equilibrated for 30min at 37 °C. Endothelial removal in 10 μM phenylephrine (PE)-pre-constricted vessels was confirmed by the loss of a vasodilatory response to 10 μM acetylcholine (Ach) at 60 mmHg. To evaluate the myogenic response, vessels were pressurized to 80 mmHg, and observed for the development of myogenic constrictions. Pressure was then dropped back to 10 mmHg. Vessels with a confirmed response were used. The vessel was then subjected to 10min pressure steps, from 10 to 120 mmHg, to allow for the development of stable myogenic tone. Following a pressure return to 10 mmHg, the NB was replaced with Ca^2+^-free NB (same constitution as NB but with no CaCl_2_ and 2 mM EGTA added). All chemicals were purchased from Sigma-Aldrich (St. Louis, MO). Data was collected as the vessel diameter at the end of the 10min at each pressure step and expressed as a percentage of the diameter change relative to the initial diameter [(D_x_ – D_10_)/D_10_*100].

### Mesenteric Artery Reactivity

Mesenteric arteries were isolated and mounted in the pressure myography system as above. Arteries were equilibrated at 37°C for 30min at 10mmHg and initial responsiveness tested with 10 μM phenylephrine (PE) and 10 μM acetylcholine chloride (Ach) at 60 mmHg. Vessels showing at least 40% dilation were used for subsequent experiments. Vessels were pre-constricted with 10mM PE and outer diameter measurements were collected after 1 min with increasing Ach or papaverine concentrations [1×10^−11^-1×10^−3^ μM] using the MyoVIEW 4 software. Increasing concentrations of PE [1×10^−11^-1×10^−3^ μM] or Angiotensin II [1×10^−11^-1×10^−5^ μM] were used to measure constriction responses. In experiments where NOS inhibition was performed, 10μM L-NAME was added to the bath superfusate; the vessel was then incubated at 10 mmHg for 30 minutes prior to Ach and papaverine responses in the presence of the inhibitor. The average of 2-3 vessels per agonist was taken for each mouse, with 6-8 mice per genotype. Phenylephrine, acetylcholine chloride, angiotensin II human, papaverine and L-NAME were purchased from Sigma-Aldrich (St. Louis, MO).

### RNA Isolation and Quantitative Real Time PCR

Following euthanasia with CO_2_, 3-month-old *Fbln4*^*+/+*^ and *Fbln4*^*E57K/E57K*^ mice were perfused with cold 1x phosphate buffered saline (PBS). The mesenteric arterial bed was dissected in cold 1x PBS and placed in RNAlater (Invitrogen, Waltham, MA) at -80ºC until mRNA was extracted using TRIzol (Invitrogen, Waltham, MA) per manufacturer’s protocol. Reverse transcription of 1ug RNA was done using the High-Capacity RNA-to-cDNA Kit while quantitative polymerase chain reaction (qPCR) was done using 50ng of cDNA, TaqMan Fast Universal PCR Master Mix, and TaqMan assays (primers/probes). Reactions were run in duplicate on QuantStudio 3 system, and experimental gene expression was normalized to that of *B2m*. TaqMan assays used in this study were Mm01208059_m1 (*Nos1*), Mm00440502_m1 (*Nos2*), Mm00435217_m1 (*Nos3*), and mm00437762_m1(*B2m*). All reagents were purchased from Applied Biosystems (Waltham, MA).

### Protein Isolation and Western Blotting

The mesenteric arterial bed of 3-month-old *Fbln4*^*+/+*^ and *Fbln4*^*E57K/E57K*^ mice were isolated as above and homogenized in 50mM Tris-HCl, pH 7.5, 1% Triton-X containing 1x phosphatase inhibitor (EMD Millipore, Burlington, MA) and 1x protease inhibitor (Sigma-Aldrich, St. Louis, MO) using a TissueLyzer (Qiagen, Germany). Protein lysates were quantified using BCA assay (Pierce, Waltham, MA) and equivalent amounts were subjected to SDS-PAGE on a 4-15% TGX polyacrylamide gel (Bio-Rad, Hercules, CA). Proteins were transferred onto a polyvinylidene difluoride membrane (Bio-Rad, Hercules, CA) overnight at 90 mA for 18hrs at 4°C. Membranes were blocked in 5% BSA/0.1% Tween20 in 1x TBS or 5% milk/0.1% Tween20 in 1x PBS for primary antibody mouse anti-human phospho-eNOS (1:1,000, pS1177, #612392, BD Biosciences, Franklin Lakes, NJ) or mouse anti-human-eNOS (1:1,000, #610297, BD Biosciences, Franklin Lakes, NJ), respectively. Membranes were then incubated with secondary antibody goat anti-mouse IgG-HRP (1:10,000, KPL). Protein was detected using SuperSignal West Atto ECL substrate reagent (ThermoFisher, Waltham, MA). Membranes were initially probed with phospho-eNOS, stripped using Restore Plus Western Stripping Buffer (Pierce, Waltham, MA) and then probed for total eNOS. Loading controls were determined using mouse anti-B-actin antibody (1:4,000, Sigma-Aldrich, St. Louis, MO), goat anti-mouse IgG-HRP (1:20,000, KPL), and detected using Clarity ECL Western Blotting Substrate (Bio-Rad, Hercules, CA). Densitometry analysis was performed using ImageJ^59^.

### Dihydroethidium Stating

Dihydroethidium (DHE) staining was performed following the protocol described in ^60^. Briefly, after euthanasia and thoracotomy, the right atrium was clipped, and 5 mL cold (4°C) 1x PBS was flushed through the left ventricle to clear the blood. The gut was removed *en bloc* and third order mesenteric arteries were dissected on ice. Vessels were placed into NB buffer for 5 min at 37°C and then incubated with 10uM DHE (Invitrogen, Waltham, MA) in NB buffer at 37°C for 20min. Vessels were then fixed in 4% paraformaldehyde for 10 min and buffer containing DAPI (1:2,000, Thermofisher, Waltham, MA) for 5 min. Vessels were mounted with ProLong Gold Antifade mountant (Thermofisher, Waltham, MA) and coverslipped. Z-stack images of the vessel wall were obtained using a FV1000 Olympus Confocal microscope and FV10-ASW 3.0 software at 20x. Detection settings were set the same for all images. ImageJ-Fiji software^59^ was used to create a maximum intensity image of the z-stack and a 52μmx71μm area of the vessel was defined as the region of interest (ROI) where integrated pixel intensity (mean intensity/area) was quantified. Data from three ROI per mesenteric artery were averaged for each animal. The entire procedure (from mouse euthanasia to image acquisition) occurred within 8 hours.

### Golgi Immunostaining and Cell Polarity Determination

Third order mesenteric arteries were collected as described above. Briefly, arteries were excised with a small section of the 2^nd^ order arterial branch for orientation of blood flow direction. After fixation with 4% paraformaldehyde at 4°C overnight, vessels were incubated in 1x PBS and cut open longitudinally. Vessels were then incubated in blocking/permeabilization buffer: 3% BSA, 1% fish gelatin, 0.5% Triton-X in 1x PBS for 1 hr, primary anti-Golgi antibody (1:100, GM130, BD Biociences) at 4°C overnight, followed by secondary antibody, goat anti-mouse Alexa Fluor-594 (1:1,000, ThermoFisher) for 1hr at RT. DAPI (1:2,000, Thermofisher) was used for nuclear labeling. Vessels were mounted on a glass slide with ProLong Gold antifade reagent (Thermofisher, Waltham, MA) and cured overnight. Z-stack images of the endothelial cell layer were collected as described above with a confocal microscope and 60x objective. A maximum intensity image of the endothelial cells was used to determine polarity and the angle between the vector of the flow direction (as determined by 2^nd^ order branch location) and the vector of the Golgi (line from center of nucleus to the center of Golgi) was measured using Image J-Fiji^59^. Cells were classified as with the flow having an angle 0º-30º, sided as between 30º and 155º, and against the flow as 156º-180º.

### Alexa-633 Hydrazide Staining and Fenestrae Number Quantification

Following euthanasia, third order mesenteric arteries were excised from 3-month-old *Fbln4*^*+/+*^ and *Fbln4*^*E57K/E57K*^ mice that had been flushed with PBS, fixed with 10% formalin buffered solution (VWR) and stored at 4°C until ready to use. Vessels were transferred into PBS and cut open longitudinally. Subsequent incubations were done at room temperature. After incubating the vessels in blocking buffer: 1%BSA (Sigma), 1% fish gelatin (Sigma) in 1x PBS for 30min, they were transferred to a solution of DAPI (1:2,000, ThermoFisher) in blocking buffer for 30min. The solution was replaced with 0.4uM Alexa633-hydrazide (ThermoFisher) in blocking buffer and vessels were incubated for 10min. Vessels were then mounted on a glass slide with vectamount permanent mounting medium (Vector Laboratories, Burlingame, CA). Z-stack images were obtained with a FV1000 Olympus Confocal microscope and FV10-ASW 3.0 software at 60x. Using ImageJ-Fiji and a LungJ plugin (Wollatz, et al., 2016, *<u>LungJ v0</u>*.*<u>5</u>*.*<u>1</u>*, University of Southampton, doi:<u>10.5258/SOTON/401280)</u>, a maximum intensity image was created from the z-stack and the number of holes manually counted. Statistical analysis was done taking the average of 3 different regions of the arteries per mouse.

### Statistical Analysis

Student’s t-test, one-way or two-way analysis of variance with multiple comparisons test was used to determine differences between genotypes, as indicated in each figure legend. Statistical analyses were run using Prism 9 for Mac OS X (GraphPad Software Inc.). Differences were considered statistically significant when *P* was equal to or less than 0.05.

## Acknowledgments

This work was supported by National Institutes of Health grants K08-HL135400 to CMH and R01-HL53325 to RPM. Funds were also provided by the Ines Mandl Research Foundation to RPM and Washington University Pediatric Nephrology Research Core to CMH.

## Author Contributions

C.M.H. and R.P.M. conceived of the study. C.M.H., M.L., and R.P.M. designed and interpreted the experiments. M.L. performed all the experiments. K.J and B.M.B. maintained the mouse colony and performed genotyping. C.M.H. drafted the manuscript, which was reviewed, edited, and approved by all authors.

